# Comparative evaluation of full-length isoform quantification from RNA-Seq

**DOI:** 10.1101/698605

**Authors:** Dimitra Sarantopoulou, Thomas G. Brooks, Soumyashant Nayak, Anthonijo Mrcela, Nicholas F. Lahens, Gregory R. Grant

## Abstract

Full-length isoform quantification from RNA-Seq is a key goal in transcriptomics analyses and has been an area of active development since the beginning. The fundamental difficulty stems from the fact that RNA transcripts are long, while RNA-Seq reads are short. Here we use simulated benchmarking data that reflects many properties of real data, including polymorphisms, intron signal and non-uniform coverage, allowing for systematic comparative analyses of isoform quantification accuracy and its impact on differential expression analysis. Genome, transcriptome and pseudo alignment-based methods are included; and a simple approach is included as a baseline control. Salmon, kallisto, RSEM, and Cufflinks exhibit the highest accuracy on idealized data, while on more realistic data they do not perform dramatically better than the simple approach. We determine the structural parameters with the greatest impact on quantification accuracy to be length and sequence compression complexity and not so much the number of isoforms. The effect of incomplete annotation on performance is also investigated. Overall, the tested methods show sufficient divergence from the truth to suggest that full-length isoform quantification and isoform level DE should still be employed selectively.

## Background

Alternative splicing and isoform switching play central roles in cell function; and disruption of the splicing mechanism is associated with many diseases and drug targets (1,2). The function of a protein is ultimately determined by its full complement of functional domains. Differential splicing typically involves a reshuffling of the functional domains to construct a functionally different protein. Gene level analyses must ignore these differences. Before things like pathway enrichment analysis can be brought down to the transcript level, it will be necessary to quantify expression of full-length isoforms.

For investigations specifically focused on splicing, one also has the option of working at the local splicing level (e.g., MAJIQ(7)). If, for example, full-length isoform quantification simply leads to an exon skipping event, that would have also been found by local splicing methods. Investigators must therefore carefully factor in the goals of their analysis to decide at which level features should be quantified.

Another reason for estimating isoform level expression is to give better estimates of gene level expression. Indeed, it is not clear how to achieve gene level quantification from local splicing information. For various purposes, full length isoform quantification must be more informative than local splicing information when it can be achieved, and the primary reason local splicing methods are popular right now is due to the relative difficulty in working with full length.

The fact that isoform quantification is a key goal for modern transcriptomic profiling is reflected in how active the community has been in developing methods and how popular those methods have been, in spite of their notoriously high false positive rates. Despite many published algorithms, in practice effective quantification of full-length isoforms from short-read RNA-Seq remains problematic and therefore has never been routine. The fundamental limitation is that individual short reads do not contain information on long-range interactions that would associate splicing events that are separated by more than the fragment length. Regardless, methods can exploit additional biological and stochastic information, like canonical splice sites, which combined with alignment information can increase accuracy (3–6).

Although long sequence read technology is improving, compared to short read technology it continues to be lower throughput with a much higher base-wise error rate and is generally more expensive. Therefore, most RNA-Seq studies are still performed with short reads and this will likely remain the case until competing technologies mature. Short reads are typically 100-150 bases long, and usually obtained from both ends of short 200-500 base fragments. Meanwhile a significant portion of RNA transcripts are over 1000 bases and many are much longer.

Given the difficulty in full-length isoform quantification, many RNA-Seq studies simply quantify at the gene level, which is much easier because uniquely aligning reads are rarely ambiguous at the gene level. Indeed, unless the investigator is specifically interested in splicing, gene level analysis will likely lead to the same conclusions, since all isoforms of the same gene typically have the same pathway annotations.

Meaningful unbiased benchmarking conclusions rely on independent investigations and realistic benchmarking data where the ground truth is known or well-approximated. There are in fact a few independent studies that compare the performance of transcript quantification methods using simulated data (8), real data (9), or a hybrid approach with both real and simulated data (10–12). So why did we embark on another comparative study? Angelini *et al* (8) and Kanitz *et al* (12) are five and six years old, respectively, and hence they do not reflect the recent developments in this fast-changing field. For instance, they do not include the popular pseudo-alignment-based methods *kallisto* (19) and *Salmon* (18). Angelini *et al* (8) take the approach of using simulated data, which is most similar to the approach employed here, however, they utilize the FLUX simulator which does not allow for many of the effects of real data we can model using the BEERS simulator (16). Also, the primary focus is on detection of isoforms as being “present/absent” in the sample and accuracy of quantification was presented as tables of quantiles. Their conclusion was that “all tables indicate that the problem of obtaining reliable estimates is still open.” Therefore, these methods require ongoing evaluation by the user community.

Zhang *et al* (10) use the human universal reference sample (UHRR) and the human brain universal reference (HBRR) which are such artificial samples that it is not clear what practical guidance can be drawn from the results. In particular the UHRR is a mixture of 10 cancer cell lines. Cancer transcriptomes are notoriously scrambled and mutated, and therefore represent a very special case, particularly with regards to annotation-based quantification. Moreover, a mixture of ten such cell lines give a sample so different from what researchers use in practice that it precludes the possibility of evaluating the methods in the context of a typical differential expression analysis, which is the main goal of most RNA-Seq studies. With the UHRR and HBRR samples only technical replicates can be generated, while all DE methods require biological replicates. Simulated data which mimic real samples is arguably more realistic than real data obtained from mixtures of 10 cancer cell lines. *In silico* simulated data offer more control as the truth is known exactly, but these data invariably simplify some of the inherent complexities of real data. In 2016, Teng *et al* (13) published very nice guidance on quantification benchmarking. Their approach assumes one has benchmarking data where the truth is known on the level of differential expression, without assuming as known the actual quantified values. Since the goal here is to investigate quantification accuracy directly, the methods in Teng *et al* are not directly applicable. Other studies focus only on single-cell data (14), or on differential splicing (15). Commonly, RNA-Seq transcript level quantification is validated by PCR. However, PCR is low-throughput and is based on probes that interrogate only a small part of a given transcript; it is also sensitive to biases at the amplification step. On the other hand, in *in silico* simulated data the truth is known exactly.

*Hayer et al* (11) investigated *de novo* transcriptome assembly, where isoform structures need to be inferred directly from the RNA-Seq data and concluded that none of the evaluated methods is accurate enough for routine use and further method development is required. The problem we investigate is considerably easier; isoform level annotation is given and reads must just be assigned to the correct isoform.

Approaches for quantifying isoform expression can be divided into three main categories. The first approach uses reads mapped to the genome by an intron-aware aligner, e.g. STAR (17). The genome alignment information is then used to assign quantified values to transcripts (3–6). The second approach is similar to the first, except it is based on reads aligned directly to the transcriptome, rather than the genome (6,17). The third approach follows the concept of pseudo-alignment which prioritizes execution performance and does not involve *bona fide* alignment (18,19). In reality, all genome aligners are transcriptome-aware, and transcriptome alignments are genome aware, so the distinction is not as cut and dried as it once was. But nonetheless, we continue to distinguish the two, with caveats. There are many published methods for quantifying full-length isoforms, however the vast majority of studies performing isoform specific analysis have used Cufflinks, RSEM or some simple counting method following genome alignment (Fig 1) (20–22). Pseudo-aligners were introduced more recently and therefore have lower adoption but are beginning to see wider usage (23,24).

**Fig 1.**
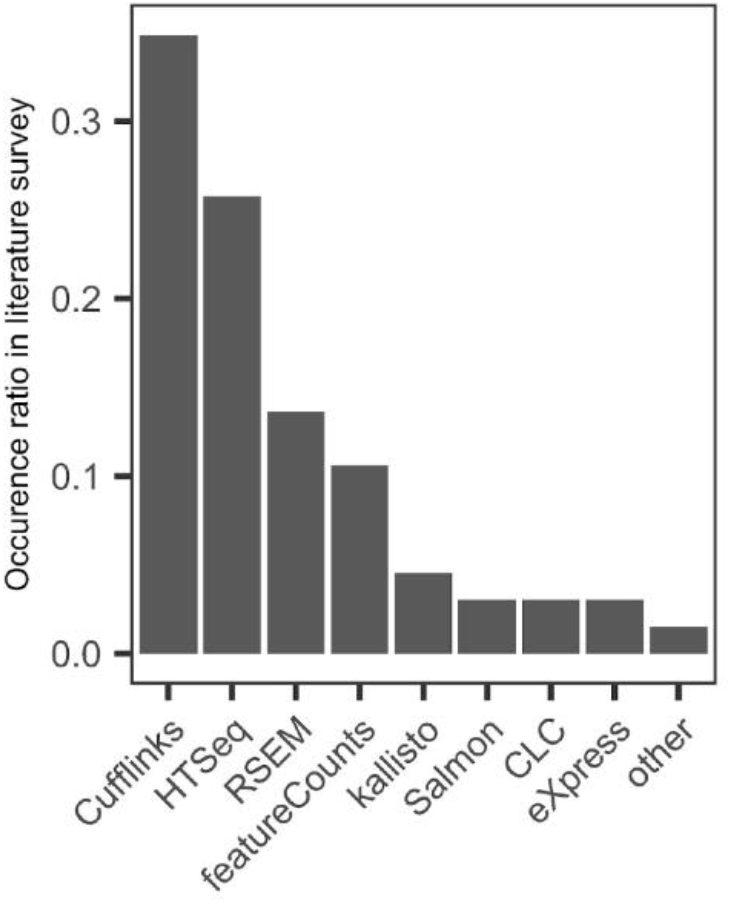
Most popular quantification methods. Ranking of quantification methods by the number of times found in the 100 most recent RNA-Seq studies (published

Here we present a benchmarking analysis of the six most popular isoform quantification methods: kallisto, Salmon, RSEM, Cufflinks, HTSeq, and featureCounts, based on a survey of the literature (Fig 1). HTSeq and featureCounts are not recommended by the authors for full-length isoform quantification, however they were included for the purpose of comparison and because they are used in practice. We also include a naïve read proportioning method, based on employing the distribution of signal inferred from the unambiguous read alignments to portion out the ambiguous read alignments, similar to the method first described by Mortazavi *et al* (40). We generated datasets from two mouse tissues, Liver and Hippocampus, which are known to be quite different in terms of splicing, with brain generally being more complex than any other tissue.

A hybrid approach is taken to obtaining benchmarking data, where real samples are emulated to generate simulated data where the true isoform abundances are known; this was done using a modified version of the BEERS simulator (16). Idealized data were generated to obtain upper bounds on the accuracy of all methods. Data were also generated with variants, sequencing errors, intron signal and non-uniform coverage, to assess how they affect performance. Since annotation is never perfect, we evaluate performance while varying annotation completeness.

Usually, the aim of an RNA-Seq analysis is to inform a downstream differential expression (DE) analysis. Therefore, we also evaluate the methods on this level, using both real and simulated data. However, it is much more challenging to produce realistic data with known ground truth at the DE level. Unlike isoform level quantification which is sample-specific, DE ground truth is established at the population level, and therefore involves much more complex benchmarking data. Our simulated samples reflect the complex joint distribution of expression across biological replicates, and thus it is meaningful to perform a DE analysis on them. This is described in more detail below but briefly, in lieu of knowing the ground truth in terms of which isoforms are differentially expressed, for each method we compare the DE analysis performed on the known true isoform quantifications of the simulated data to the DE analyses performed on the estimated counts determined using the particular method. The more different the two analyses are, the less accurate the quantification method must be in informing the DE analysis. This then allows us to compare the methods in terms of their accuracy of quantification. It is possible that a method underperforms another method at the level of quantification, but outperforms it in the DE analysis.

## Results

### Hybrid benchmarking study using both real and simulated data

For the simulated data we started with 11 real RNA-Seq samples: six liver and six hippocampus samples from the Mouse Genome Project (25). Isoform expression distributions were estimated from these samples in (7) which were then used to generate simulated data for which the source isoform of every read is known. Two types of simulated datasets were generated with the BEERS simulator (16). First, idealized simulated data were generated, with no SNPs, indels, or sequencing errors, no intron signal and uniform coverage across each isoform (7). Second, simulated data were generated with polymorphisms (SNPs and indels), sequencing errors, intron signal, and empirically inferred non-uniform coverage (7). Relative performance on idealized data does not necessarily reflect relative performance on real data, but we do expect the accuracy of the methods on idealized data to be upper bounds on the accuracy in practice. If a bound on idealized data is below what one would tolerate in practice, then it cannot be expected to be viable in practice. The (more) realistic data provide insight into the effect of the various factors on the method performance. The realistic data probably also gives bounds on accuracy of real data since it was designed to be no more complex than real data. For simplicity of exposition, we will refer to the data with the complexities as the “realistic” data, keeping in mind it does not reflect every property of real data, just the five properties listed above (SNPs, Indels, Sequencing Error, Intron Signal and Non-Uniform Coverage). For both the idealized and realistic simulated data, we use three liver and three hippocampus samples to evaluate isoform quantification, and six liver and five hippocampus samples to evaluate DE analysis, as in [21]. All samples were obtained from independent animals raised as biological replicates. Comparisons between tissues were employed to assess consistency and differential expression; brain has a more complex transcriptome than other tissues (26), and thus isoform level analysis is expected to be more challenging for the algorithms.

We performed a comparative analysis of seven of the most commonly used full-length isoform quantification algorithms; kallisto (19), Salmon (18), RSEM (6), Cufflinks (4), HTSeq (3), featureCounts (5) and a naïve read proportioning approach similar to the method first described by Mortazavi *et al* (40) (NRP; See Methods). Kallisto and Salmon are pseudo-aligners; RSEM, Cufflinks, HTSeq, and featureCounts are genome alignment-based approaches where the alignments are guided by incorporating transcriptome information, and NRP is a transcriptome alignment-based approach. These methods were evaluated at the isoform expression level using idealized and realistic simulated data, with full and incomplete annotation, and also at the differential expression level using both realistic and real data.

### Comparison of full-length quantification methods

#### Idealized data

The idealized data has no indels, SNP’s, or errors, includes no intron signal, and deviates from uniform coverage across each isoform only as much as may happen due to random sampling. Under such perfect conditions we expect that all methods will achieve their best performance. The data were aligned to the reference genome or transcriptome with STAR (17) and quantified with the seven methods. In Fig 2A, estimated expression is plotted against the true transcript counts, for each method in Liver. Each point represents the average of the three replicates of that tissue. A point on the diagonal indicates a perfect estimate. A point on the *X*-axis indicates an unexpressed transcript which was erroneously given positive expression. A point on the *Y*-axis indicates an expressed transcript which was erroneously given zero expression.

**Fig 2.**
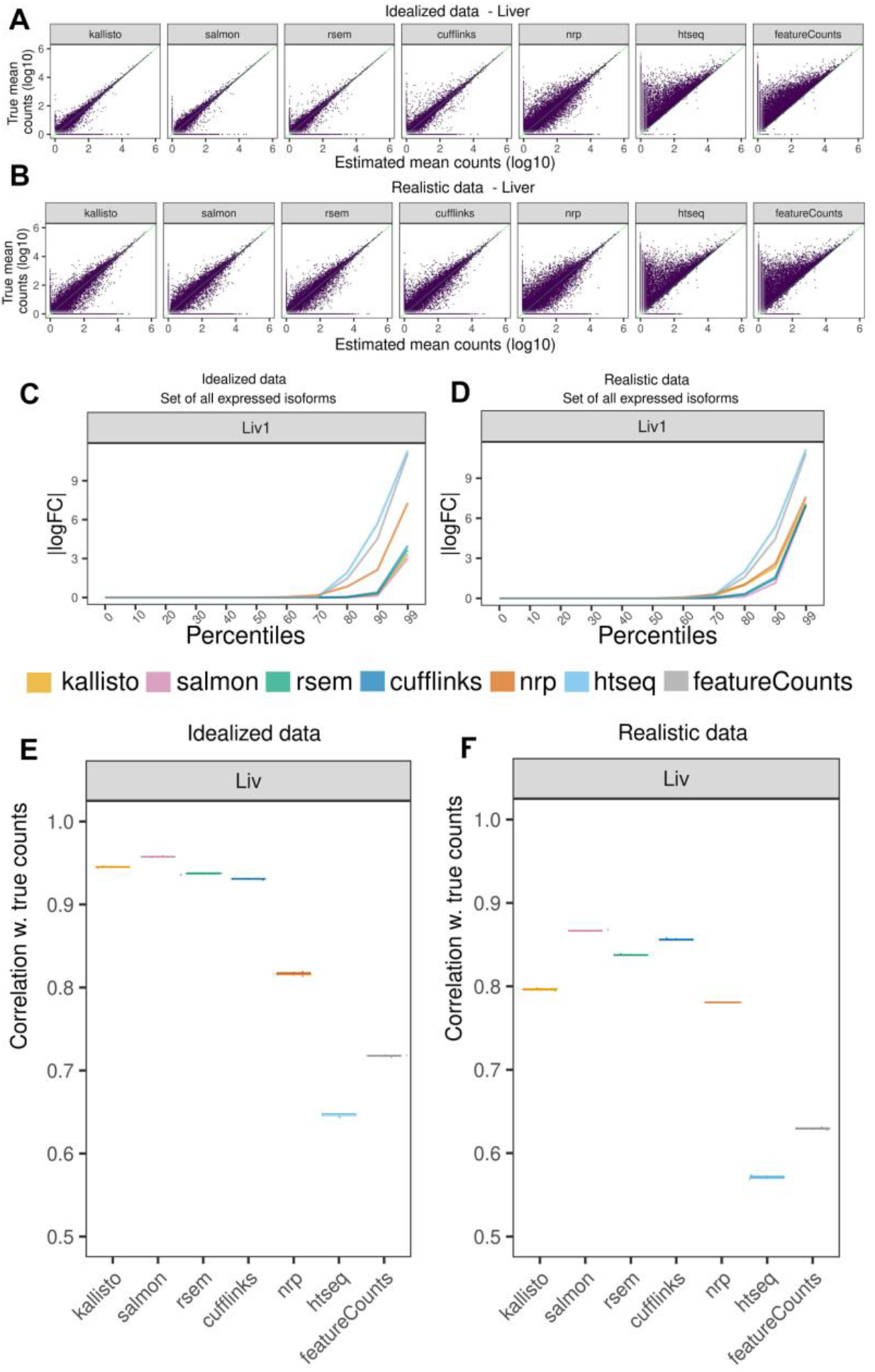
Comparison of estimated quantification with the truth in simulated data. (A,B) Scatter plots between the inferred and true counts. Each point represents the average expression of three samples. A) idealized data B) realistic data. (C,D) Percentiles of the |logFC| (relative to true counts), for the set of expressed isoforms in sample 1 in C) idealized and D) realistic data. A point (*x,y*) on a graph means *x*% of the transcripts have |logFC|≤*y*. For clarity, all unexpressed genes were removed. Unexpressed isoforms were retained if a sibling isoform is expressed. A logFC of 0 means the estimated count is equal to the truth for that isoform. (E,F) A point on this plot shows the Spearman correlation between the vector of inferred quants and the truth. For each method, the correlation was calculated for each of the three samples, plotted as a box plot. Variation is low, so box plots are thin.

In liver, we observe that kallisto, Salmon, RSEM and Cufflinks’ are fairly concordant with the truth, while NRP performs moderately well considering its simplicity. HTSeq and featureCounts show high deviation from the truth and in particular tend to undercount. That is because both methods ignore any read which is ambiguous between isoforms.

Accuracy is somewhat lower in hippocampus, which has more complicated transcription with more alternate splicing, but similar conclusions hold (Fig S1A) for all methods.

In what follows we will regularly use the log (base 2) fold change between the inferred counts relative to the truth. This will be denoted logFC and if the absolute value is taken then |logFC|. Before taking ratios, all counts are adjusted by addition of a pseudo-count of 1 (see Methods).

Scatter plots are only so informative. Fig 2C and S1C compare for each method the percent of transcripts that have |logFC|*<c* for each *c>*0. Specifically, a point (*x,y*) on the graph means *x*% of transcripts have |logFC|<*y*. The lines are monotonically non-decreasing and the closer to the *x*-axis, the better. When the |logFC| for a transcript is close to zero, it suggests that the method gives an accurate estimate. When the *p*-th percentile of the cumulative distribution is close to zero it means the top *p* percent of the best estimated transcripts are all very accurate. Genes where all of their isoforms have zero expression will be generally easy for the algorithms to get right, since there will usually be no reads mapping anywhere to the gene locus. To avoid those exaggerating the average performance, such genes were removed from consideration before generating these graphs. These graphs were also generated with all genes removed that had three or less expressed isoforms (Fig S1E), to focus just on the more difficult cases, which visually makes very little difference to these accuracy percentiles. Kallisto, Salmon, RSEM, and Cufflinks display high accuracy up to the 85th percentile. NRP diverges from the truth at the 60th percentile. We see minimal difference between replicates or between tissues (Fig. S1C).

For each sample, correlation was computed between the vectors of true and inferred quants (Fig 2E, S1 Table). All four methods kallisto, Salmon, RSEM and Cufflinks perform comparably, with Salmon marginally better. One must keep in mind rank correlation is a very general measure which could mean different things, so it is important interpret with caution. But nonetheless the considerably lower correlation in NRP, HTSeq and featureCounts is notable.

#### Realistic data

The “realistic” data include variants (SNPs and indels), sequencing errors, intron signal, and non-uniform coverage across each isoform. Non-uniform coverage mimics observed coverage in real data but does not attempt to model coverage-associated sequence features such as GC content. As such, the non-uniform coverage options of Salmon that involve GC content could not be utilized. The non-uniform coverage option of RSEM was used since it is sequence agnostic.

Similar to the idealized data, points in Fig 2B and S1B are the average of the three quantified versus the average of three true counts, for each tissue. In both tissues, kallisto, Salmon, RSEM, Cufflinks, and NRP’s isoform quantification is still largely concentrated around the diagonal (Fig 2B, S1B), but the variance from the truth is markedly increased, as compared to the variance in the idealized data. NRP’s performance on realistic data has decreased less relative to the idealized than other methods, so the difference between methods is now less apparent. Similar to the results with the idealized data, HTSeq and featureCounts consistently undercount across datasets.

The percentile plots of |logFC| are given in Figs 2D, S1D. These show that Salmon, RSEM and Cufflinks continue to be highly concordant with the truth up to the 80th percentile. NRP and kallisto now perform comparably. HTSeq and featureCounts diverge from the truth the fastest. Comparing the percentile plots and scatter plots, it appears the error profile of the most accurate 80% are roughly the same in the realistic as the idealized, but the least accurate 20% are considerably less accurate in the realistic than the idealized. We will investigate below in more detail what features of the isoforms are driving the accuracy, but it is not just the number of isoforms. Length plays perhaps a larger role. And other features come into play as well, such as sequence compression complexity.

The correlation boxplots for the realistic data are given in Fig 2F (S2 Table). Here we see a greater separation between the methods. Salmon has incurred relatively less of an increase in error in the realistic data as compared to kallisto, indicating there is more difference between the two than there was in early versions, where the primary difference was in how GC bias was handled. This data has no GC bias, so that cannot explain the difference here. Surprisingly, the accuracy of NRP has barely been reduced in the realistic data and is comparable to kallisto, indicating this approach may have some robustness that the other methods lack and could perhaps be modified to be competitive.

Salmon is marginally superior here, in both absolute comparisons and correlations.

### Hierarchical Relationships Between the Methods

Clustering was performed to investigate the hierarchical relationships between the methods. Here, the number of replicates was increased to be all six liver and all five hippocampus realistic samples. The hierarchical clustering was based on the average expression, with correlation-based distance metric. Hippocampus and Liver give somewhat different results (Fig 3A-B). The true counts cluster with the first group. Surprisingly, Cufflinks goes from a neighbor of the truth in hippocampus to an outlier in liver. Meanwhile, clustering on the logFC of hippocampus *versus* liver, shows all methods reasonably close to the truth except HTSeq and featureCounts (Fig 3C).

**Fig 3.**
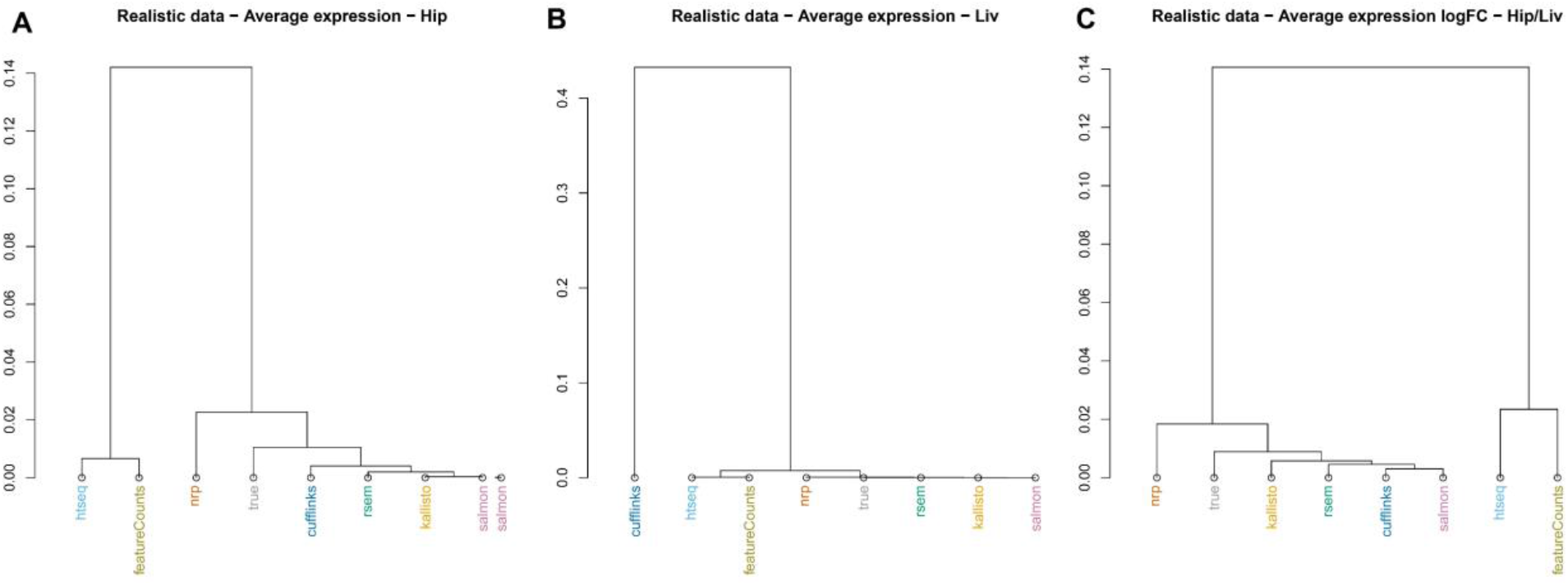
Similarity of quantification methods. Using realistic data (5 hippocampus and 6 liver samples). Hierarchical clustering with correlation distance, on average expression, for A) hippocampus and B) liver. C) Hierarchical clustering with correlation distance, on average expression |logFC| of hippocampus over liver.

### Features associated with quantification accuracy

We next investigate the covariates that affect the quantification accuracy. For example, the more isoforms a gene has, the more difficult we expect the problem to be. Other obvious features that we expect to impact accuracy are length and number of exons. Less obvious features will be investigated below, but first we look at length. The logFC of estimated counts relative to true counts is plotted against transcript length in Fig 4A-B and S2A-B. Only transcripts of length 200 bp or longer were simulated, so the plots start there. Surprisingly, the variance from the truth does not increase monotonically with length as much as might be expected. All methods appear to have the most difficulty with moderate length isoforms. A value of ±10 on the vertical axis represents a fold-change from the truth of roughly 1000-fold, so regardless of length there is considerable divergence from the truth for all methods.

**Fig 4.**
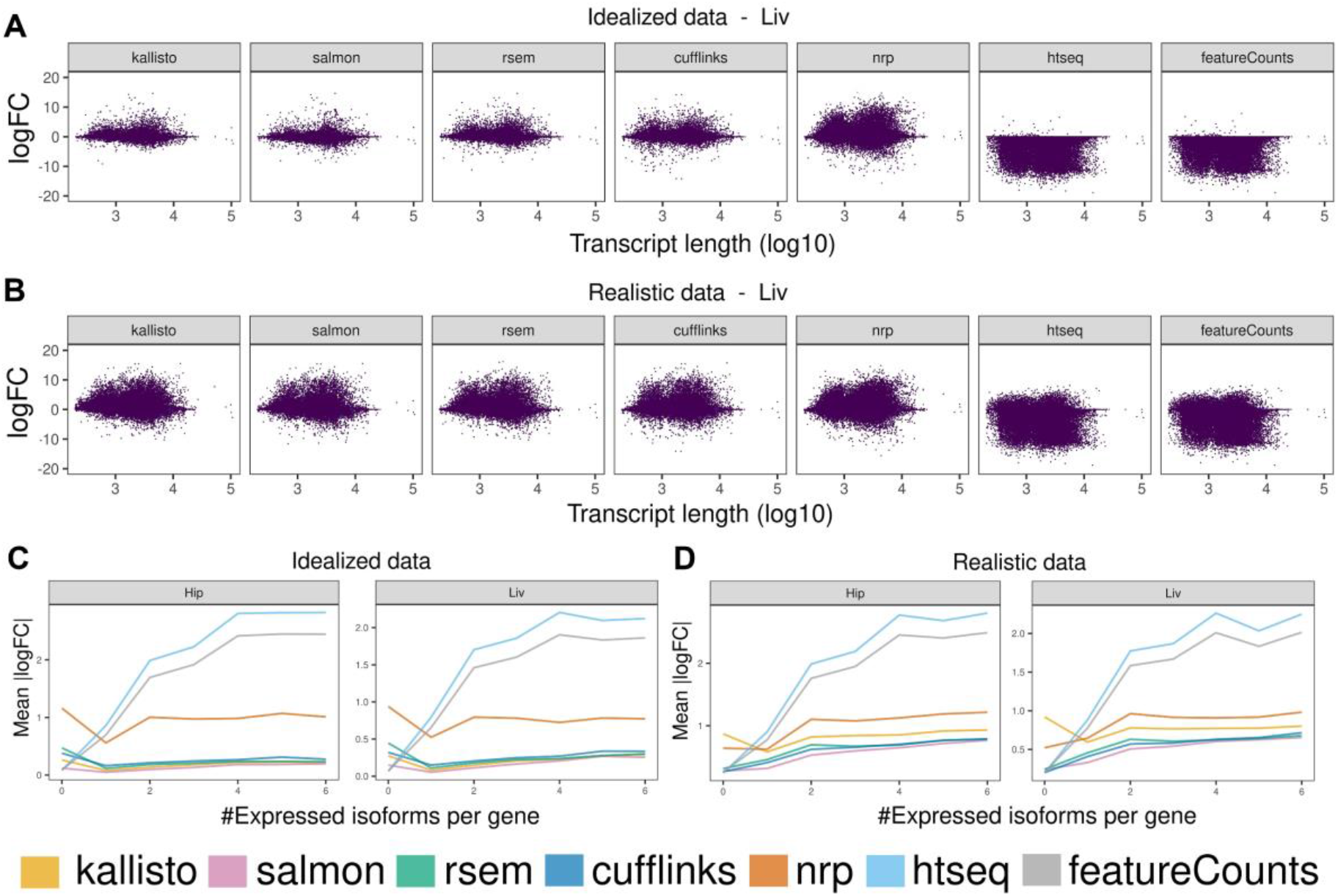
Effects on quantification accuracy. Effect of transcript length on the concordance of each method to the truth, given the average of three liver samples, using a) idealized and b) realistic data. Effect of number of expressed isoforms on the mean |logFC| for both tissues, using c) idealized and d) realistic data. The maximum number of expressed isoforms between the three replicates, and the mean of the |logFC|s at a given number of expressed isoforms are plotted. The mean |logFC| is based on at least 100 transcripts.

Next Fig 4C-D shows the |logFC| plotted against the number of isoforms (S3-4 Table). The closer to zero the |logFC| is, the better. Surprisingly, mean accuracy of the viable methods kallisto, Salmon, RSEM and Cufflinks does not decay rapidly with the number of expressed isoforms. In realistic data, the number of isoforms effect is larger.

Additionally, to investigate whether the methods are more accurate with genes with low number of isoforms, we filtered out the genes with 1 or 2 isoforms and saw that the performance of the most accurate methods (kallisto, Salmon, RSEM, Cufflinks, NRP) is largely unaffected (Fig S1E-F).

Another way to investigate the features associated with accuracy is to compare the distribution of the feature over all isoforms, to the distribution of the feature over the set of isoforms most discordant from the true counts. For convenience these will be called the “background” and the “foreground” distributions. Specifically, in each tissue in the realistic data, for each method, the 2,000 transcripts were identified with the greatest |logFC| relative to the truth (averaged across three of the replicates). This produces, for each method/tissue, a list of the most discordant isoforms. For a given structural property, such as length, the background distribution is the (empirical) distribution of the property over all isoforms (which is the same for both tissues). And the foreground distribution is the (empirical) distribution of the property over the 2,000 discordant isoforms. These two distributions can be plotted together and interpreted visually - and can also be compared using the Kolmogorov-Smirnov test. The Teqila tool was used for this analysis (28).

A number of structural properties were investigated, such as number of isoforms, hexamer entropy, transcript length, transcript sequence compression complexity (32), exon count, etc. The *p*-values were multiple-testing corrected by Bonferroni for testing multiple methods and multiple properties. The significant properties are shown in Fig 5, the more significant the darker. In the idealized data, exon count and transcript length are comparable. However, in the realistic data length becomes much more relevant. Surprisingly, the number of sibling isoforms (other isoforms of the same gene) is far less relevant than the length or number of exons, except for NRP where the number of sibling isoforms is strongly associated with accuracy.

**Fig 5.**
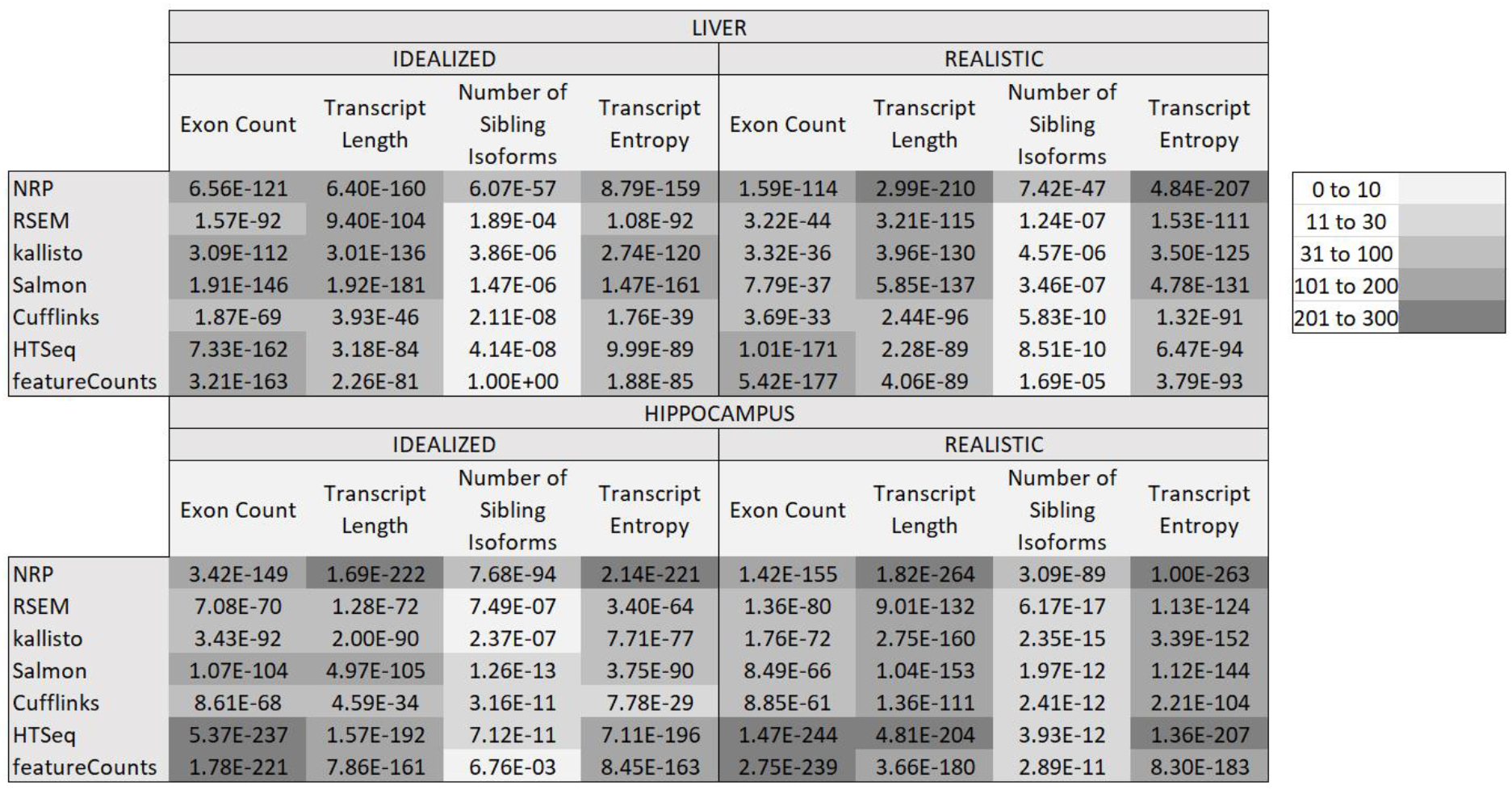
Features Associated with Inaccuracy. This table shows the Kolmogorov-Smirnov (multiple-testing corrected) *p*-values between the distribution of the metric over all isoforms, and its distribution over the 2000 most discordant isoforms, determined by the |logFC| to the truth. In the realistic data the transcript length and transcript sequence entropy are the most highly associated. Surprisingly, the number of isoforms is considerably less important.

It is worth looking at the distributions of some of the properties. A select few are shown here, others are in Supplemental. Fig 6 shows the foreground and background distributions for Transcript Length and Exon Count, for Salmon. Only Salmon is shown because the distributions are similar for all methods. Fig 6A shows that there are proportionally far fewer isoforms in the foreground than the background up to length of around 1000 bases. Then after a length of about 2000 bases there are proportionally far more isoforms in the foreground; however, it does not then get progressively worse. Similarly, with the number of exons, performance seems to flip at around 10 exons. With sequence compression complexity the foreground distribution is highly enriched for low compression complexity, for all methods (See Fig S3).

**Fig 6.**
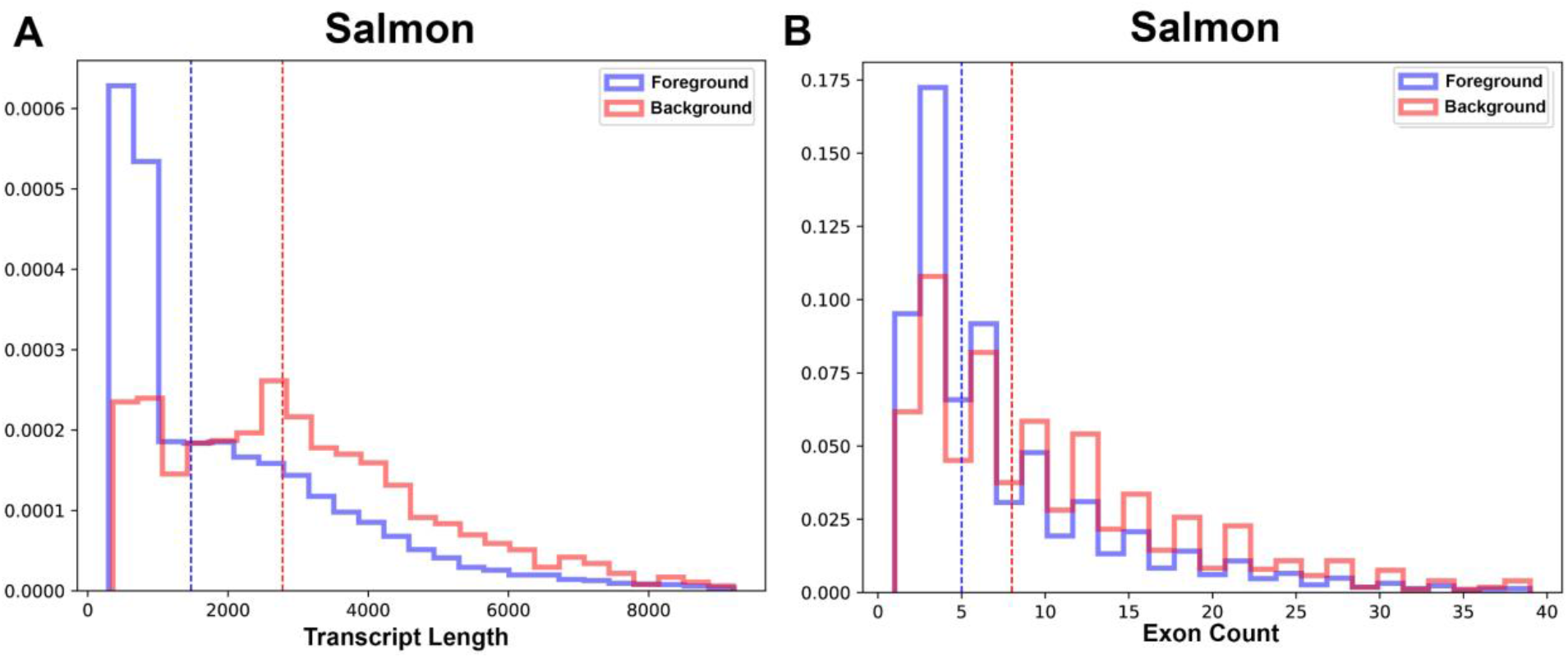
Features Associated with Inaccuracy: Transcript Length and Exon Count. Foreground and Background distributions for (A) Transcript Length and (B) Exon Count for Salmon (all methods are similar) for the realistic Hippocampus data. The sawtooth pattern in the Exon Count is just an artifact of binning, both curves should be strictly decreasing. Length starts to become problematic around 2000 bases, while exon count starts to be problematic around 10 exons. Only Salmon is shown because the distributions are similar for all methods.

The observations about length and exon count apply equally well to all methods. However, for number of isoforms, NRP is quite different from the others (see Fig 7).

**Fig 7.**
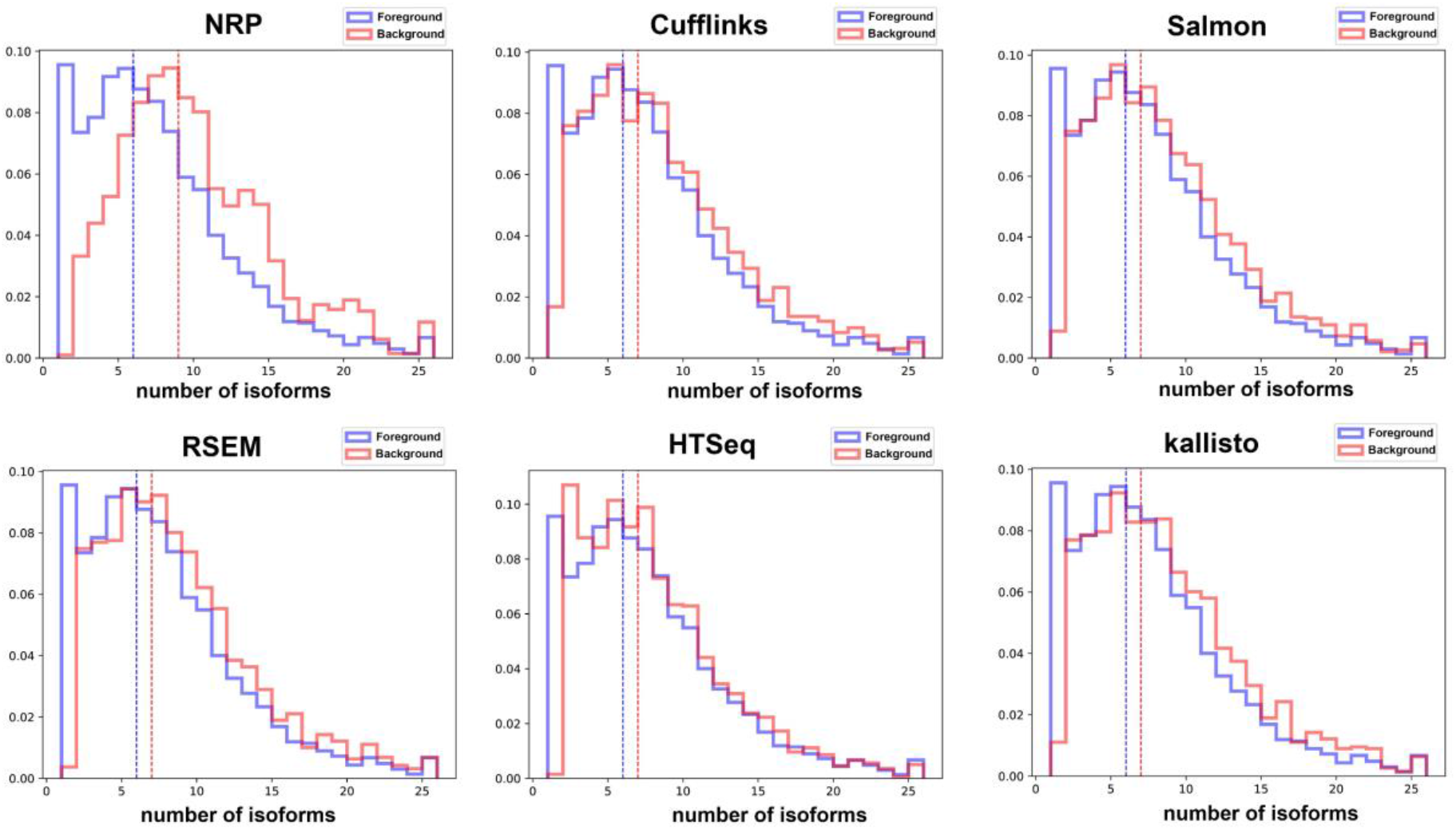
Features Associated with Inaccuracy: Number of Isoforms. Foreground and Background distributions for Number of Sibling Isoforms, for the realistic hippocampus data. All methods are shown except featureCounts which is nearly identical to HTSeq. Surprisingly, for all methods except NRP accuracy does not progressively decrease with the number of isoforms, except for the one-isoform genes, where the foreground distribution is dramatically depleted. NRP on the other hand presents a foreground distribution that is more intuitive.

### Incomplete Annotation

Annotation guided quantification is only as good as the annotation itself. And no annotation is perfect, they all contain isoforms that do not exist and are missing isoforms that do exist. It can quickly get complicated trying to sort out the effect of annotation issues on the performance of the algorithms. Sometimes missing annotation can be beneficial, for example if a non-expressed isoform is missing from the annotation, then it cannot erroneously be assigned reads. On the other hand, if a highly expressed isoform is missing, then the method must figure out what to do with the orphaned reads. It should be able to figure out that they should be ignored. Otherwise, they will be assigned to the wrong isoform. To investigate these two issues, we modified the annotation in two ways. First, all isoforms that were not expressed were removed. The extent to which their absence improves accuracy gives an indication of how well the method is handling unexpressed isoforms when they are present. Second, the highest expressed isoform of each gene was removed. This should cause major problems to the algorithms, so it is informative to see which methods handle this better. For most methods changing the annotation requires going all the way back to the alignment step.

Fig 8 shows the percentile plots of the |logFC|. Hippocampus sample Hip1 is shown but all samples Hip and Liv look very similar. The first thing to note is that removing the maximally expressed isoform has dramatically decreased the accuracy of all methods except for HTSeq and featureCounts. And removing the non-expressed isoforms has marginally increased accuracy for those methods. In contrast, for HTSeq and featureCounts we observe the opposite. Removing the non-expressed isoforms has dramatically decreased accuracy and removing the highest expressed isoform has made very little difference, particularly with featureCounts.

**Fig 8.**
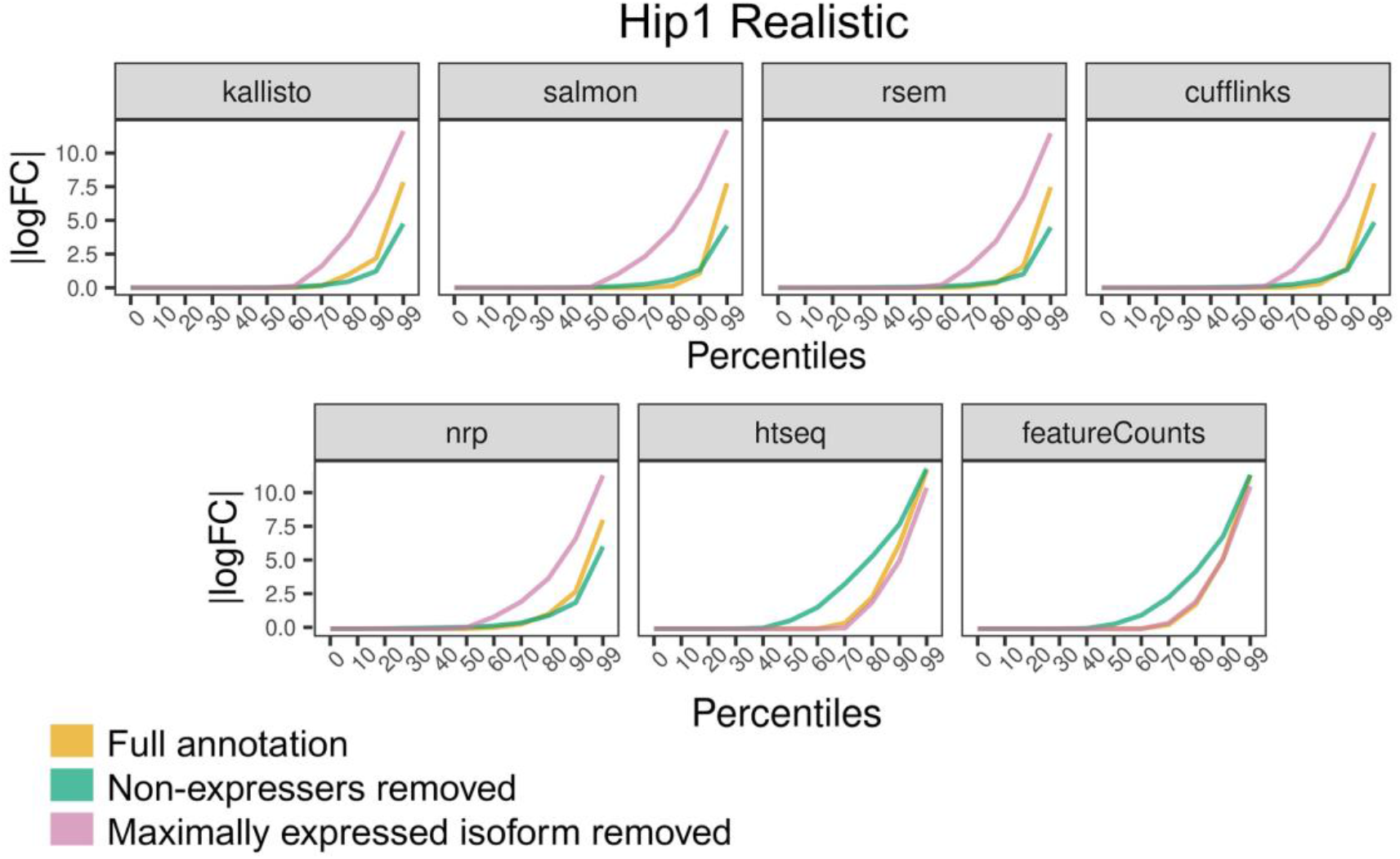
Incomplete Annotation. The annotation was modified in two ways. First, all unexpressed isoforms were removed, which should make things easier on the algorithms. Second, the highest expressed isoform of every gene was removed, which should make things much more difficult. For each method the percentile plots are shown. Here a point (*x,y*) on a curve means *x*% of isoforms have |logFC|>*y*.

Fig 9 compares for the different methods the percentile plots for removing the maximally expressed isoform. This eliminates the isoform of the majority of the reads so should have a dramatic effect on accuracy. Here Salmon has the most difficulty and HTSeq and featureCounts are the most robust to this, followed by NRP. Here we see a significant difference between Salmon and kallisto that goes in the opposite direction of the differences seen by the other perspectives.

**Fig 9.**
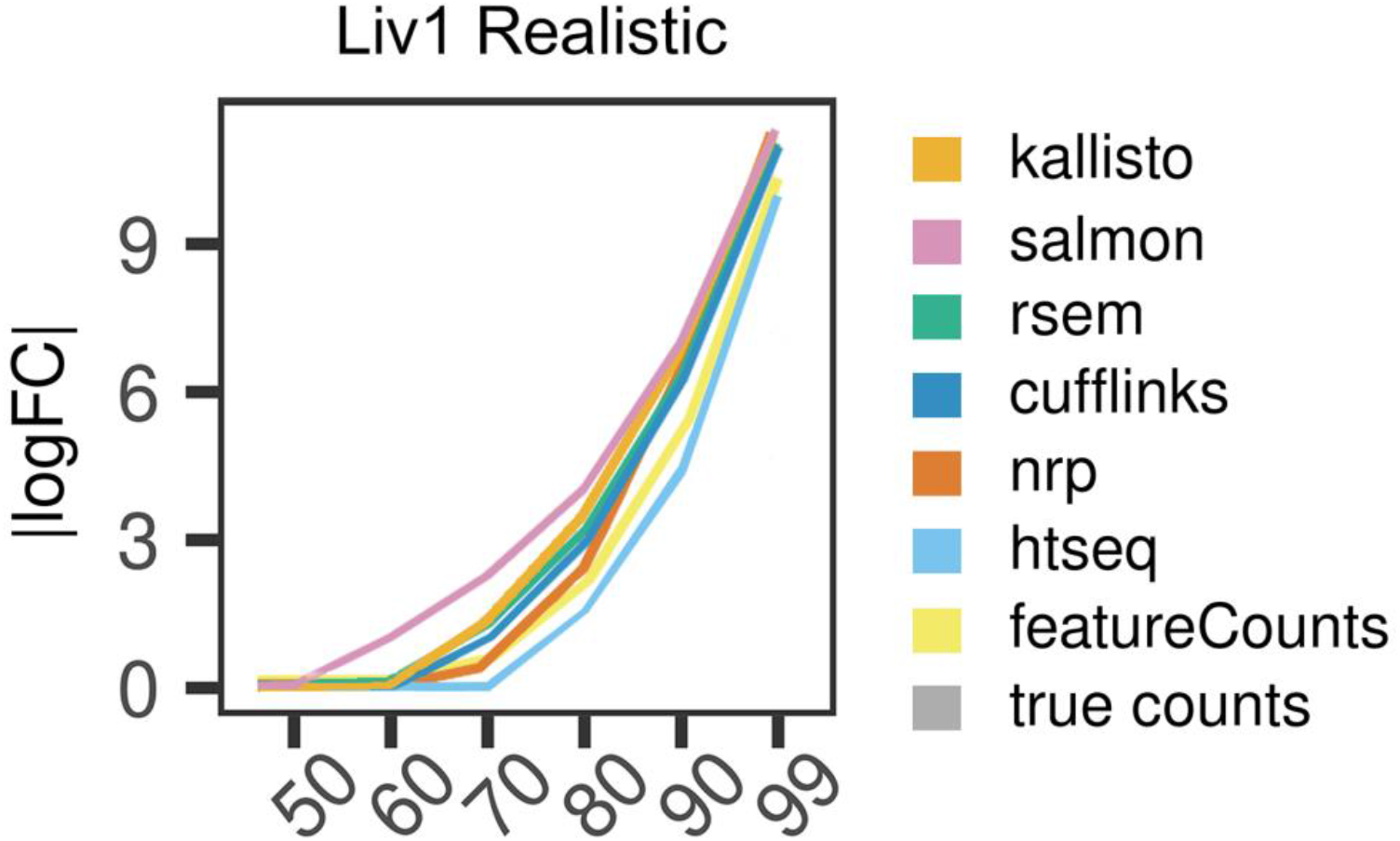
Removal of highest expressed isoforms. The annotation was modified by removing the highest expressed isoform of every gene. For each method the percentile plots are shown. Here a point (*x,y*) on a curve means *x*% of isoforms have |logFC|>*y*. The lower the curve, the better. Surprisingly, Salmon has the most difficulty and HTSeq the least.

### Effects on differential expression

Next, we use differential expression to assess quantification accuracy. If differential expression analysis is the downstream goal for the quantified values, then it does not matter if the absolute abundances differ from the truth, if the DE *p*-values are unaffected. To investigate this, the two tissues were compared against each other; different enough tissues so that there is an abundance of differentially expressed genes. Six hippocampus samples and six liver samples of the realistic data were quantified, with each of the seven methods, and the resulting quantified values were used as input for DE analyses with EBSeq (41), which is optimized for isoform differential expression. The *p*-values generated from the true counts are compared to *p*-values from the inferred counts - the assumption being that the closer a DE analysis on the inferred counts is to the corresponding DE analysis on the true counts, the more effectively the method has quantified the expression, *with respect to informing the DE analysis*. Kallisto and Salmon are recommended to be run with Sleuth, however since Sleuth cannot take true counts as input, the comparison would not be meaningful. Since we are comparing EBSeq (truth) to EBSeq (inferred) for all methods, it should be meaningful to compare methods to each other with this metric.

Comparing two developmentally divergent tissues, we expect the majority of transcripts that are expressed to be differentially expressed. Figure 10A shows the overlap with the truth, for the top *n* most significant genes, as *n* varies from 1 to 50,000. Since EBSeq reports a lot of zero *p*-values, rounded down from their limit of precision, ties were broken with the logFC. The vertical axis is the Jaccard index (29) of the top *n* DE transcripts determined using the real counts and the top *n* DE transcripts determined using the inferred counts. The Jaccard index of two subsets of a set is the size of the intersection divided by the size of the union. The higher the curve, the better. Salmon and Cufflinks are performing best from this perspective, followed by RSEM. NRP and kallisto appear roughly equivalent. In Fig 10B the number of DE transcripts is plotted as a function of the *q*-value cutoff (S5 Table). If a curve rises above the truth, then that method must be reporting more false-positives than the *q*-value indicates. At varying places between 0.05 and 0.2 all methods become anti-conservative. Salmon and Cufflinks track the truth closest at small cutoffs.

**Fig 10.**
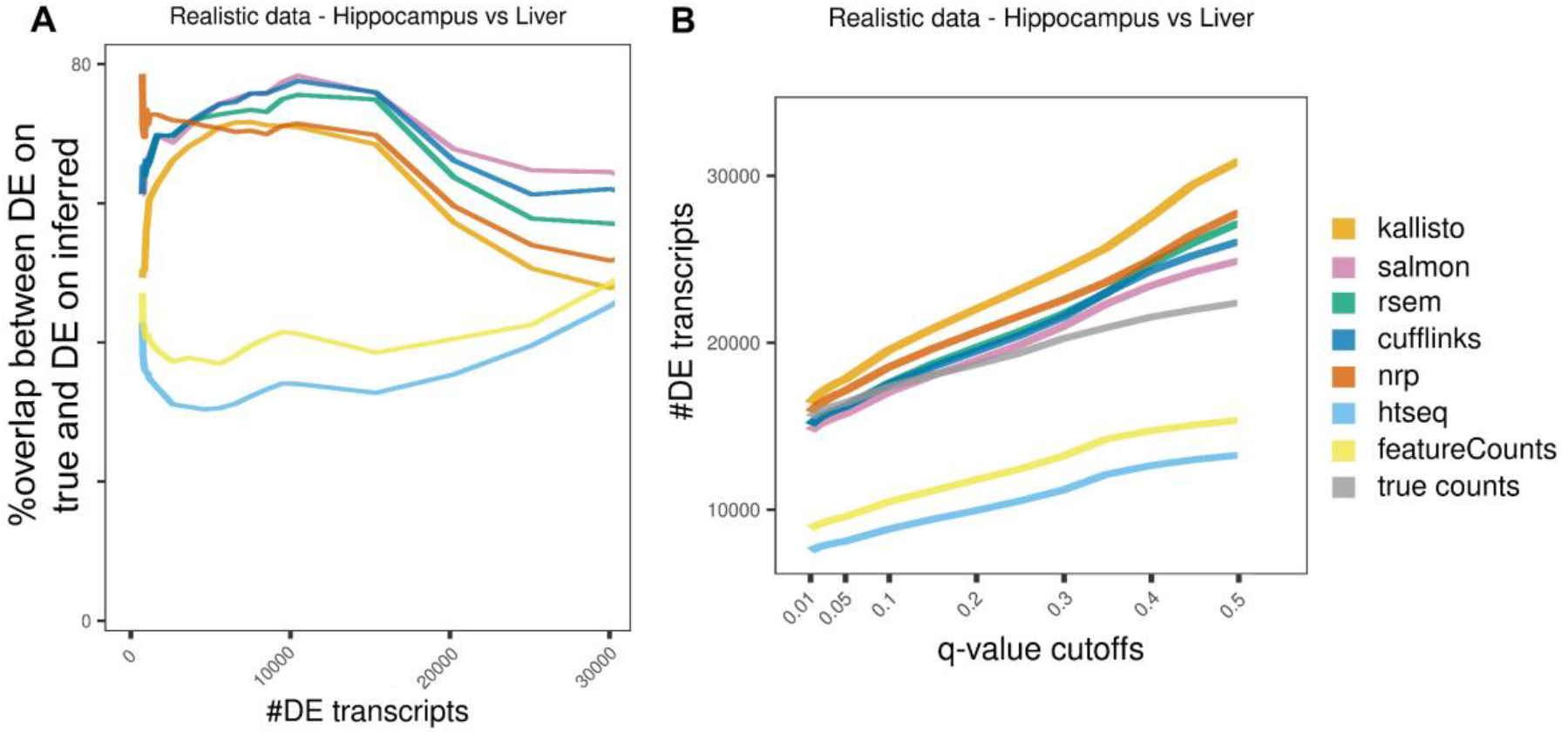
Method effect on differential expression analysis, using realistic data. For each method, a DE analysis with EBSeq was performed between the two tissues. (A) A point (*x,y*) on a curve means for the top *x* DE transcripts using real counts, and the top *x* DE transcripts using the inferred quants, have Jaccard index *y*. (B) A point (*x,y*) on a curve means there are *y* isoforms with *q*-value ≤ *x*. The curves should be evaluated in relation to the truth, which is the gray curve. At varying *q*-value cutoffs between 0.05 and 0.2 all methods become anti-conservative. Salmon and Cufflinks track the truth closest at small cutoffs.

This data can also be used to evaluate the DE methods themselves – EBSeq, Sleuth and DESeq2. DESeq is included for reference, but it was not specifically designed with transcript-level DE in mind. In DE benchmarking, it is notoriously difficult to determine a benchmark set of either differential, or non-differential, transcripts. However, if an isoform has zero expression in all replicates of both conditions, then it must necessarily be non-differential. A total of 43,872 isoforms have zero in all replicates of both conditions. Any transcript called DE in this set must be a false positive arising from mistakes in the quantification process. This allows us to define a lower bound on the actual FDR, because it gives a lower bound on the number of false positives, as given by the number of these null isoforms that were called DE. This lower bound on the FDR is plotted as a function of the *q*-value cutoff (Fig 11A). Additionally, the actual number of null isoforms called DE is plotted as a function of the *q*-value cutoff in Fig 11B. Fig 11A shows that in all cases the true FDR is much greater than reported.

**Fig 11.**
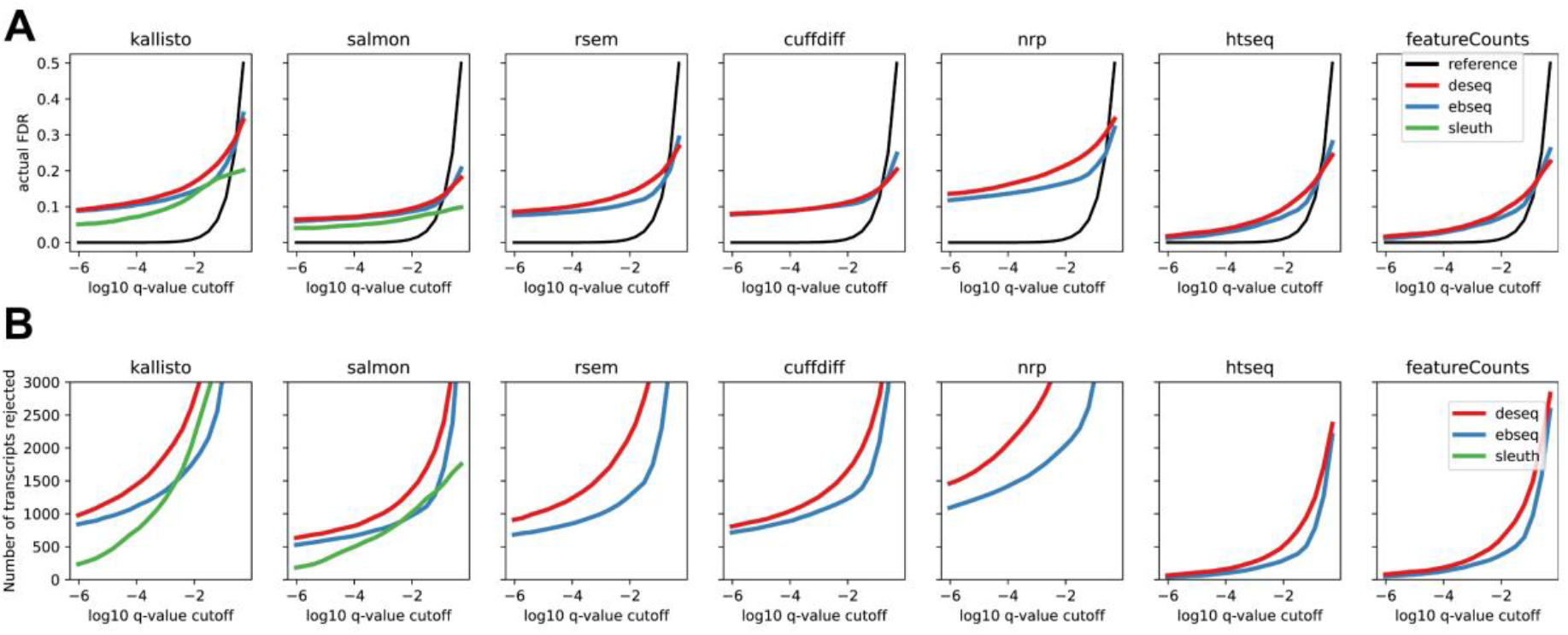
Method effect on differential expression analysis, using realistic data. The roughly 43,872 isoforms with zero true expression in both Liver and Hippocampus, serve as a set of null isoforms for the DE analysis. (A) gives a lower bound on the true FDR of the isoforms rejected at each *q*-value cutoff. Plots above the black line are anti-conservative. (B) same as A but shows the actual number of null isoforms determined DE as a function of the *q*-value. Note that only 94,929 isoforms exist in total.

Indeed, Fig 11B shows that even at very small *q*-values EBSeq and DESeq are reporting thousands of these false positives. At an FDR of 0.01 there are at least 1,000 isoforms using any method. These cannot simply be the 1% false positives allowed by an FDR of 0.01 since that would then require an additional 99,000 true positives, which is more isoforms than are even annotated.

Why is this happening? When an isoform has zero true expression, but another isoform of the same gene has positive expression, it is easy for reads of the expressed isoform to be misassigned to unexpressed. However, if none of the isoforms of a gene are expressed, it is far less likely that any of the isoforms are assigned spurious reads since it is much less likely that any reads map anywhere to the gene’s locus. Therefore, if a gene has no expressed isoforms in Liver and has one or more expressed isoform in Hippocampus, in addition to one or more unexpressed isoforms, then the unexpressed isoforms will tend to have zero expression in Liver and will tend to incur spurious expression in Hippocampus. Such isoforms are then easily mistaken as differential. An isoform level DE method should account for this variability, but we see in Fig 11 that both EBSeq and Sleuth are anti-conservative. The isoform-level DE methods do however outperform DESeq2, which is not intended for transcript-level analysis. On the quantification methods where it is applicable, Sleuth shows the lowest false positive rate, reflecting the fact that it uses additional variance information from bootstrap samples.

### Evaluation with real data

In all comparisons performed with the simulated datasets, HTSeq and featureCounts are very similar and kallisto, Salmon, RSEM, Cufflinks, and NRP are also generally comparable. To explore whether the comparative analyses can be replicated with a real experiment, we used the real data that informed the simulations. Here we used six Hippocampus and six Liver samples. Hierarchical clustering was performed with correlation distance, on the average expression of six samples. The results recapitulate these two groups in hippocampus (Fig 12A), while in liver Cufflinks clusters further and alone (Fig 12B), as in the realistic simulated data (Fig 3A-B). This suggests that Cufflinks is strongly influenced by a tissue-specific effect and confirms that the simulated data successfully capture properties of the real data.

**Fig 12.**
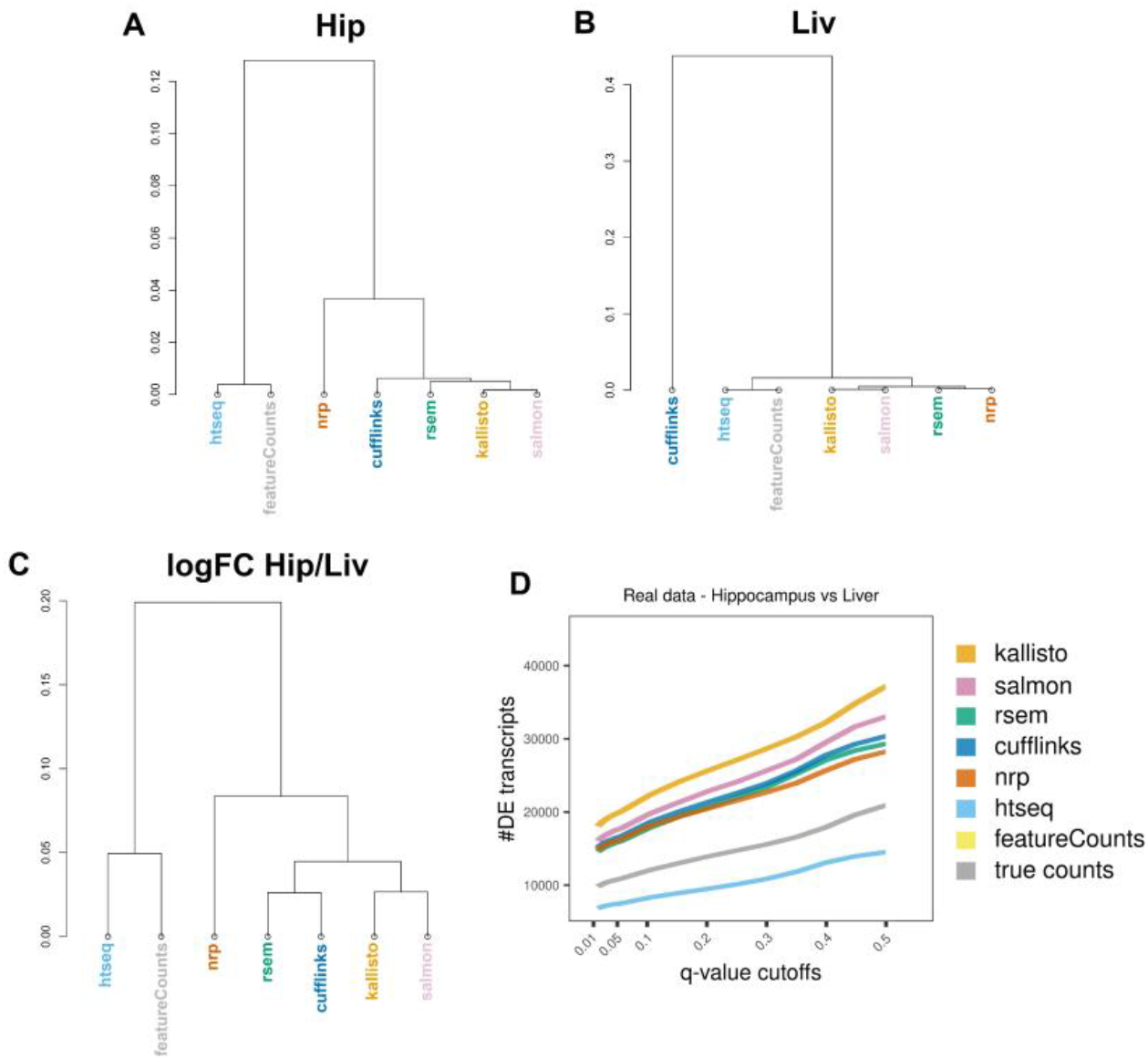
Method effect on DE analysis, using real data. Hierarchical clustering by correlation distance of the average expression using A) six liver samples or B) six hippocampus samples. C) Hierarchical clustering by correlation distance of the logFC of hippocampus over liver samples. For each method, we performed a DE analysis between the two tissues. D) Number of DE transcripts identified at various *q*-value cutoffs.

Furthermore, we compare the seven quantification approaches on how well they inform a DE analysis, using the real data. We quantified six samples from each tissue with the seven methods, followed by DE analysis between the two tissues using EBSeq. The methods cluster similarly for both realistic and real data (Figs 3,12). There is a significant difference in the number of DE transcripts identified at various *q*-value cutoffs, among the seven methods (Fig 12D, S6 Table).

## Discussion

Isoform level quantification has been an area of active development since the inception of RNA-Seq. It got off to a rough start and progressed slowly, however steadily, and we see considerable improvement over the last five years. Nevertheless, using both realistic simulated and real data, no method achieved high enough accuracy across the board that it can be recommended for general purposes.

Overall, Salmon marginally outperformed the other methods by our benchmarks. It must be kept in mind however that the additional complexities of real data will likely affect those marginal differences in unpredictable ways. Therefore, if one is going to do full length isoform quantification at this stage, then Salmon or RSEM could be equally effective choices. Cufflinks performs well from many perspectives but the erratic behavior in the Liver clustering (Fig 3,12) is concerning. Salmon, as a pseudo-aligner, has the advantage of efficiency. However, if one is performing small or medium sized RNA-Seq studies, then genome alignments should in principle always be performed anyway so that coverage plots can be examined in a genome browser. Since there is no shortcut to that process, the advantages of Salmon and kallisto in terms of efficiency really only come into play when hundreds or thousands of samples must be processed. Since data sets with hundreds of thousands of samples are on the horizon, this is a real concern. But for most targeted RNA-Seq analyses, as is done routinely in research labs, this will factor less into the decision.

Salmon (18) is similar to kallisto, and originally was identical except for incorporating a sample-specific model of fragment GC bias to improve its quantification estimates. Our simulated data, generated by BEERS (16), do not reflect these biases, and thus this feature of Salmon could not be reasonably evaluated in this study. The only simulator currently available that models fragment GC biases is Polyester (33). However, both Polyester and Salmon use the same underlying model for fragment GC bias (34), which may bias results towards Salmon’s benefit. Salmon further has options to control for read start sequence bias (such as from random hexamer priming) and positional bias (such as 5’ or 3’ bias), which were also not evaluated here. Future benchmarking studies will require datasets (both real and simulated) that capture the true sequence properties underlying non-uniform coverage in order to quantitatively assess the performance impact offered by incorporating a fragment bias model. This will be accounted for in BEERS2.0.

Additionally, we investigated some extreme cases of inaccuracy, in both simulated and real data, where transcripts were estimated to be highly expressed by one method and non-expressed by the other. In the simulated data, we identified enriched genomic properties that drive the deviation of each method from the true counts. And in real data, we isolated one example of large quantification differences between methods. In this, the inclusion of a single read causes kallisto and RSEM to disagree by 137 counts to 0, and the difference resolves if that read is removed. This edge case occurred because only two reads were unambiguous to the two isoforms of a highly expressed gene. The transcript-level DE method Sleuth (31) uses bootstrap resampling to control for possibilities like this example. EBSeq uses the number of sibling isoforms as a factor in its variance computation. However, our analysis indicates these while these methods outperform DESeq2, they could still be generating too many false positives. In particular when all isoforms of a gene are unexpressed in the first condition, and one isoform is expressed in the second condition, we observe a lot of false positives on the other unexpressed isoforms of that same gene, due entirely to quantification inaccuracy.

Overall, kallisto and Salmon as alignment-free methods require less computational time while achieving similar or better accuracy compared to other methods whereas RSEM and Cufflinks perform well among the alignment-dependent methods. However, our results indicate that all tested methods should be employed selectively, especially when long transcripts with many isoforms or transcripts with low sequence complexity are the candidates of interest for the study. NRP is a straightforward and simple approach that is relatively robust to polymorphisms, non-uniform coverage and intron signal; however, it struggles with a greater number of isoforms. In any case it performs equally well or in some cases outperforms more sophisticated methods, suggesting that information extraction and inference from short RNA-Seq reads is largely saturated and future, more complex models might offer only small benefits in gene isoform quantification.

These results indicate the differing strengths of different approaches to this problem. As such, it may be possible to leverage the different methods to achieve overall greater accuracy. For example, NRP, HTSeq and featureCounts appear to do better on one-isoform genes. So, it may make sense to treat those genes separately. In any case this must continue to be an active area of research before the technology can transform transcriptomics and realize the advantages of full-length isoform quantification.

## Methods

### Data generation

We used the same method for generating simulated data as described in *Norton et al* (7). For all of the procedures described below, we used gene models from release 75 of Ensembl GRCm38 annotation, and sequence information from the GRCm38 build of the mouse genome. We used the empirical expression levels *and percent spliced included* (PSI) values across all of the Mouse Genome Project (MGP) (25) liver and hippocampus samples estimated in *Norton et al (7)*. Briefly, the samples were aligned with STAR, and gene-level counts were calculated with htseq-count. Next, ENSEMBL transcript models were used to identify local splicing variations (LSVs); loci with exon junctions that start at the same coordinate but end at different coordinates (or vice versa). Of the 41,133 annotated genes expressed in the MGP data, 3,055 were randomly selected to reflect the empirical PSI values for their associated transcripts. For this “empirical set” of genes we estimated PSI values separately for each sample by comparing the relative ratios of all junction-spanning reads that mapped to an LSV. These PSI values reflect the biological noise and real differential splicing (if any) between the two tissues. For each of the remaining genes, we simulated no differential splicing between tissues with the following procedure: 1) For a given gene with *n* spliceforms, randomly select a gene with the same number of spliceforms from the empirical set. 2) For this empirical gene, randomly select the PSI values from one MGP sample. 3) Assign these PSI values across all samples for the gene in the simulated set. 4) To add inter-sample variability, randomly add/subtract a random number (uniform from 0 - 0.025) to the PSI values in each sample, such that PSI values for the gene/sample still sum to 1. These estimated gene expression counts and PSI values, for both the empirical set and remaining set of genes, served as input into the BEERS simulator (16). For the idealized data, we used a uniform distribution for read coverage, with no intronic signal, and no sequencing errors, substitutions, or indels (parameters: -strandspecific -error 0 - subfreq 0 -indelfreq 0 -intronfreq 0.05 -fraglength 100,250,500). For the realistic data, we used a 3’ biased distribution for read coverage that was inferred empirically from previous data (35). We also added 5% intronic signal, and used a sequencing error rate of 0.5%, a substitution frequency of 0.1%, and an indel frequency of 0.01% (parameters: - strandspecific -error 0.005 -subfreq 0.001 -indelfreq 0.0001 -intronfreq 0.05 -fraglength 100,250,500). Lastly, we did not simulate novel (unannotated) splicing events in either dataset (parameter: -palt 0).

### RNA-Seq analysis

The two simulated RNA-Seq datasets were aligned to both the GRCm38 build of the mouse genome and transcriptome with STAR-v2.7.6a (17). For all transcript models we used release 75 of the Ensembl GRCm38 annotation. The breakdown of the annotation by number of spliceforms is given in S7 Fig. The raw read counts were quantified at the transcript level, using the following methods: the pseudo-aligners kallisto-v0.46.2 (19) and salmon-v1.4.0 (18), the naïve read proportioning approach (NRP: http://bioinf.itmat.upenn.edu/BEERS/bp3/) based on transcriptome alignment, as well as the genome alignment based methods RSEM (6), Cuffdiff (Cufflinks-v2.2.1) (4,36), HTSeq-v0.12.3 (3), and featureCounts (Subread-v2.0.1) (5). EBSeq-v1.30.0 (41) was used for differential analysis, both between hippocampus and liver; and also between estimated and true transcript counts. All visualizations were done with R-v3.6.1 packages (37). The command line parameters used for each tool are in S7 Table.

#### Differential Expression Analysis

Transcript-level differential expression was assessed via three methods. DESeq2-v3.12 and EBSeq-v1.30.0 were run on the inferred quantified values from all quantification methods. In addition, the Sleuth-v0.30.0 method was run on the quantifications from Salmon and kallisto, using 50 bootstrap samples and the Wasabi package (https://github.com/COMBINE-lab/wasabi) to convert Salmon to the Sleuth input format. All methods were run on the realistic simulated data and compared the five hippocampus samples to the six liver samples and on the real samples, six hippocampus versus six liver samples. For the simulated data, we also ran DESeq2 and EBSeq given the true quantified variables for comparison with the inferred quantifications.

EBSeq was configured to perform two-condition isoform-level DE with the recommended uncertainty groups of genes with 1, 2 or 3 or more transcripts. The maxround parameter was set to 25. Since EBSeq is a Bayesian method, we used the reported posterior probability of equivalent expression as the q-value of the transcript being DE (41). Since EBSeq yields many transcripts with q=0, we broke ties by using the logFC from the quantified values, when ranking genes by q-value.

#### Description of the seven quantification methods

*Kallisto* is a pseudo-aligner which uses a hash-based approach to assemble compatibility classes of transcripts for every read by mapping the read’s *k*-mers, using the transcriptome *k*-mer *de Bruijn* graphs (19). It requires few computing resources and has a fast runtime. The index was built from the transcript sequences and transcript abundances were quantified via pseudo-alignment using the index. The counts estimates in the est counts column were used in our analyses. Fifty bootstrap runs were performed for DE analysis by sleuth.

*Salmon* is a pseudo-aligner which also accounts for various biases in the data (GC content, starting sequencing bias, position-specific fragment start location bias such as a 5’ or 3’ bias) (18). Like kallisto, it has fast runtime and low resource requirements. The index is built from transcript sequences and decoy sequences of the entire genome were provided. The NumReads estimate was used in our analysis. Fifty bootstrap runs were performed for DE analysis by sleuth.

*RSEM* is a gene/isoform abundance tool for RNA-Seq data which uses a generative model for the RNA-Seq read sequencing process with parameters given by the expression level for each isoform (6,38). A set of reference transcript sequences was built using rsem-prepare-reference script based on the GRCm38 Ensemblv75 reference genome and the corresponding transcript annotation file. Then the isoform abundances were estimated using rsem-calculate-expression. For our analysis, we use the expected count in the isoform output file which contains the sum (taken over all reads) of the posterior probability that each read comes from the isoform.

To prepare input for Cufflinks, HTSeq and featureCounts, the real and simulated data were aligned to a STAR genome index built with the GRCm38 Ensemblv75 transcript annotation file.

Cuffdiff2 (36) is an algorithm of the *Cufflinks* suite (4), which estimates expression at the transcript-level and controls for variability across replicates. Because of alternative splicing in higher eukaryotes, isoforms of most genes share large numbers of exonic sequences which leads to ambiguous mapping of reads at the transcript-level. Cuffdiff2 first estimates the transcript-level fragment counts and then updates the estimate using a measure of uncertainty which captures the confidence that a given fragment is correctly assigned to the transcript that generated it (39). We provided the sorted aligned files and the appropriate annotation file to cuffdiff2 and used the isoforms.count_tracking file generated.

For *HTSeq (3)*, htseq-count was used to estimate isoform level abundances from the alignments. We used the recommended default mode which discards any ambiguously mapped reads and hence conservative in its estimate. The HTSeq documentation suggests that one should expect sub-optimal results when it is used for transcript-level estimates and recommends performing exon-level analysis instead (using DEXSeq). Nevertheless, we use it for transcript-level fragment count estimates in order to quantify its underperformance relative to the other methods.

*featureCounts (5)* is a read count program to quantify RNA-Seq (or DNA-Seq) reads in terms of any type of genomic property (such as gene, transcript, exon, etc.). It is very similar to htseq-count, with the main differences being efficient memory management and low runtime.

As a baseline comparison, we considered a *Naïve Read Proportioning (NRP)* approach as a baseline. This is essentially the method described by Mortazavi *et al* (40) but without normalizing by transcript length. NRP uses a transcriptome alignment (provided by STAR in this case) and in the first pass, computes the number of reads mapping unambiguously to each transcript. To deal with ambiguous mappers, it then takes a second pass on the alignment file. If a read maps ambiguously to a set of transcripts *𝒯* { *T*_1_, *T*_2_, … *T*_*n*_}and *c*_1_, *c*_2_, … *c*_n_ are the respective fragment counts from unambiguous mappers in the first step, it increments the fragment count of *T*_*i*_ by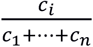. If all of the *c*_*i*_ ‘s are 0, that is, none of the transcripts in *𝒯* have any reads mapping unambiguously to them, we increment the fragment count of *T*_*i*_ by 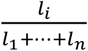 where *l*_*i*_ is the length of transcript *T*_*i*_.

### Statistical analysis

As a measure of the accuracy of each method, we compute the absolute value of the log_2_fold-change (fold-change after adjusting numerator and denominator by pseudocount of 1) for estimated counts relative to the known simulated true counts. For example, if x is the true count and y is the estimated count for a particular method, we calculate the quantity of 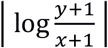 for each transcript. The closer the logFC is to 0, the more accurate the method is for that transcript.

In order to better represent the distribution of the |logFC| values for each method, we plot (for the set of expressed isoforms) the value of |logFC| corresponding to every tenth percentile starting from 0. If the method has high accuracy, we expect the graph to be close to 0. Thus, if the graph for method A is higher than method B, we conclude that tool B is more accurate.

Moreover, we identify the genomic properties of the data that affect the accuracy of the methods. For each method, we identified the most discordant transcripts sorting by |logFC|. Using the Ensembl annotation and genome sequence for GRCm38, we created a database of transcript properties (such as number of isoforms, hexamer entropy, transcript length, compression complexity* (32), exon count, etc.) and their global distributions across the transcriptome. Then for the lists of discordant transcripts, we computed the Kolmogorov-Smirnov two-sample test *p*-values for each transcript property, followed by Bonferroni correction for multiple testing, to identify the properties that exhibit significant deviation from the global distribution.

* Transcript sequence compression complexity is a metric that captures the amount of lossless compression of the transcript sequence. The higher the sequence complexity, the lower the compression, which implies higher transcript sequence compression complexity.

## List of abbreviations

logFC: log_2_ fold change
DE analysis: differential expression analysis
DE transcripts: differentially expressed transcripts
NRP: naïve read proportioning approach

## Declarations

## Acknowledgements

We thank the High Performance Computing at Penn Medicine (PMACS HPC) funded by 1S10OD012312 NIH, for the cluster computing support.

## Availability of data and materials

All raw and processed RNA-Seq data used in this study are available at Array Express under accession number E-MTAB-599. All simulated data generated in this study are available at http://bioinf.itmat.upenn.edu/BEERS/bp3/.

## Additional Files

Supplemental materials are in Sarantopoulou_FLIquant_supplemental_material.pdf

## Authors’ contributions

GG, DS, and SN conceived of and designed the study. DS, TB and SN performed all computational analysis and visualization. NL produced all RNA-Seq simulated data. All authors contributed to discussions and running the algorithms. DS, TB, SN, and GG wrote the manuscript. All authors read and approved the manuscript.

## Supplemental Figures

**S1 Fig.**
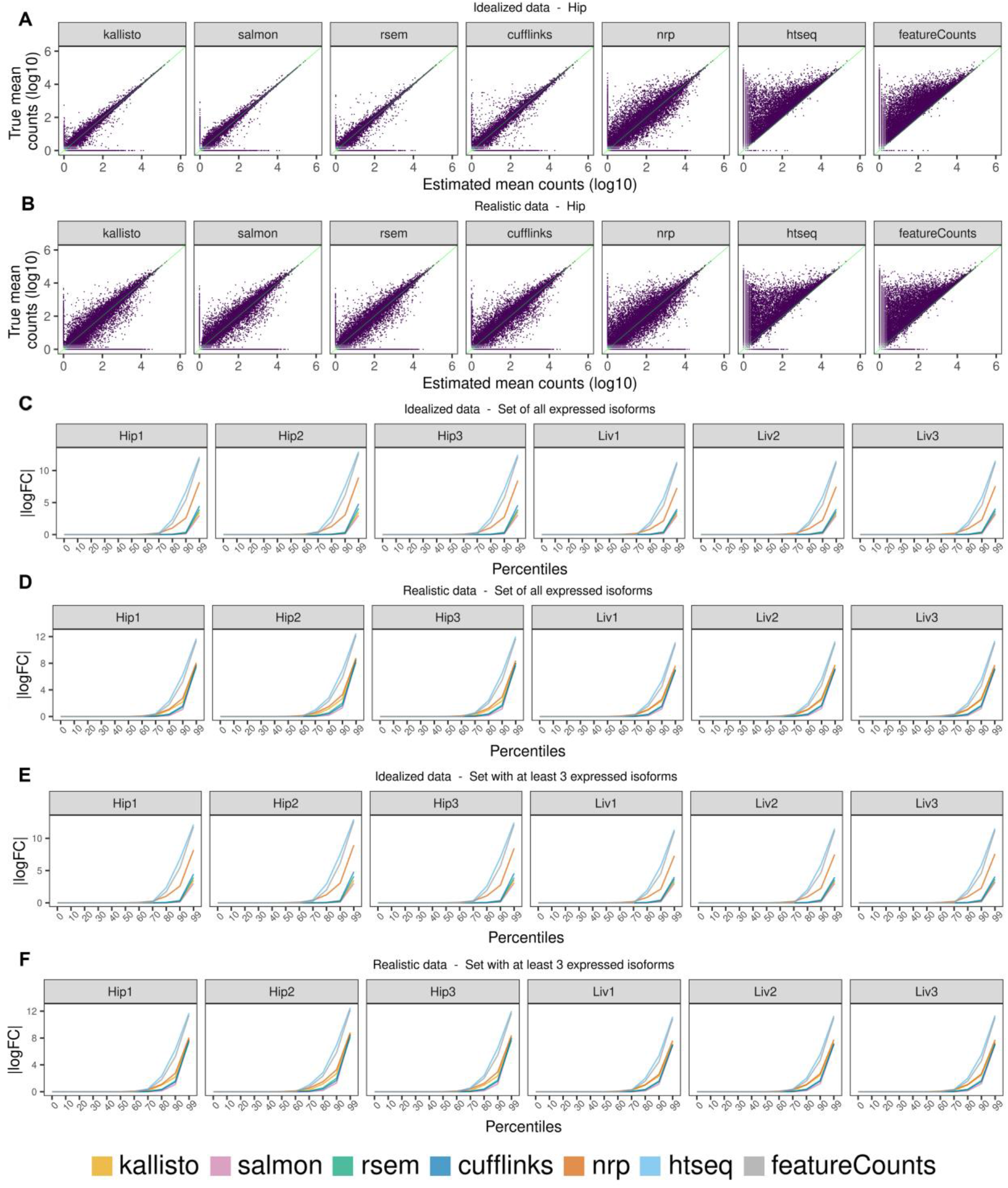
Method effect on full-length isoform quantification using simulated data. Method effect on full-length isoform quantification using simulated data. Average expression of three hippocampus samples, comparing each method to the truth, using A) idealized and B) realistic data. Percentiles of cumulative distribution of |logFC| using C) idealized data, D) realistic data, E-F) idealized and realistic data respectively, where we restricted to the set of genes that have at least 3 expressed isoforms.

**S2 Fig.**
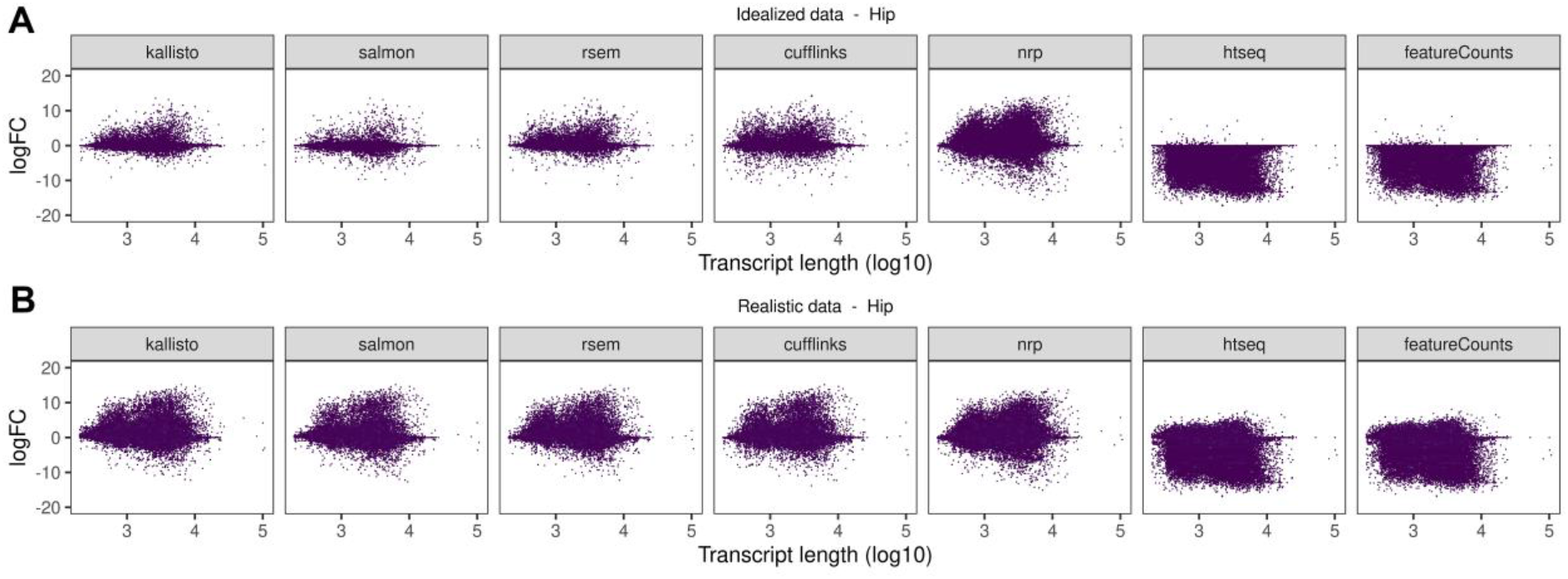
Effect of transcript length on quantification accuracy. Effect of transcript length on quantification accuracy, given by adjusted logFC of the average of the three hippocampus samples, using A) idealized and B) realistic data.

**S3 Fig.**
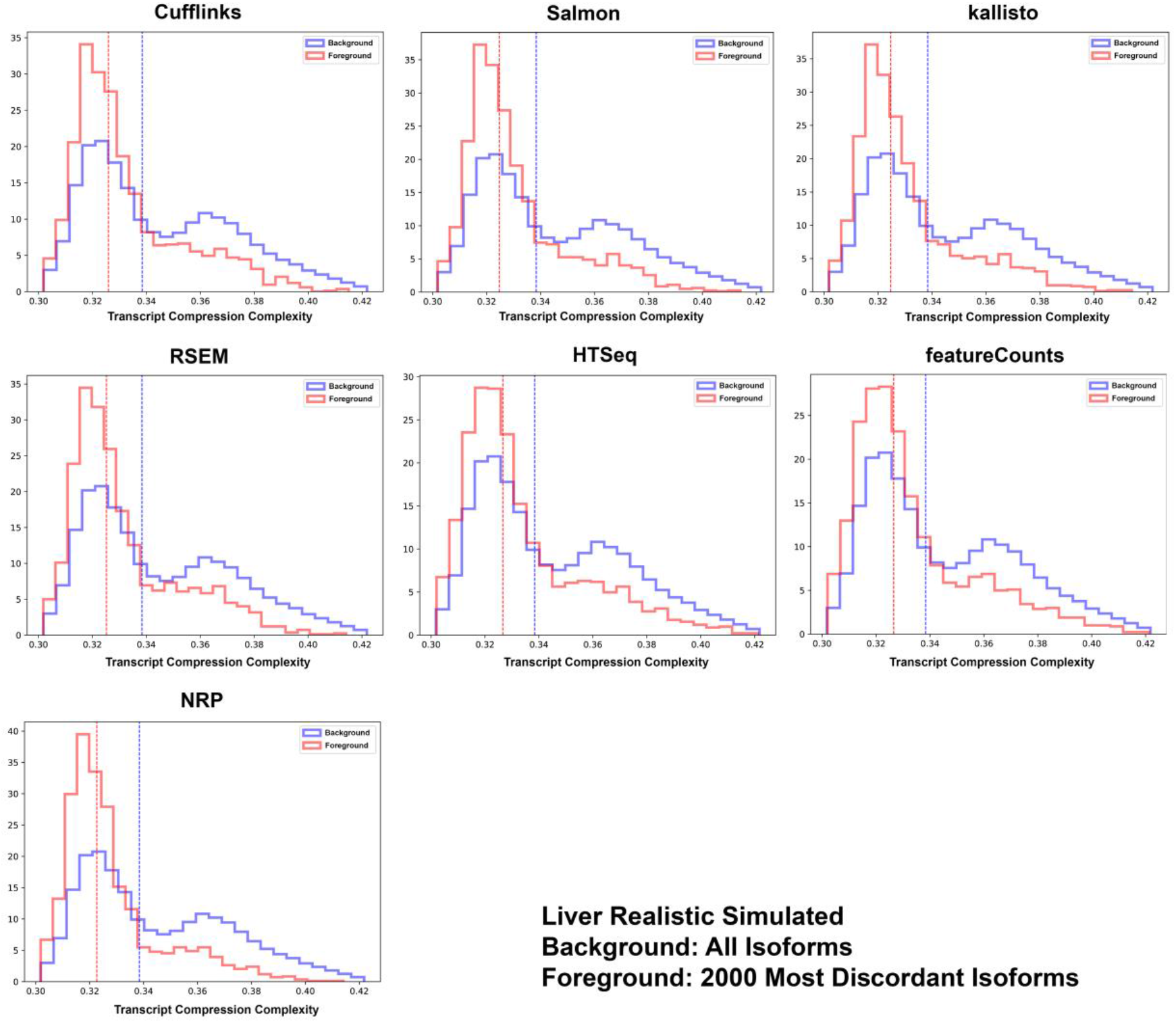
Differential distribution of transcript compression complexity. For each method the foreground and background distributions are shown for transcript compression complexity. The background is over all isoforms, the foreground is over the top 2,000 discordant transcripts sorted by absolute adjusted log2FC. The foreground distribution is highly enriched for low compression complexity for all methods.

**S4 Fig.**
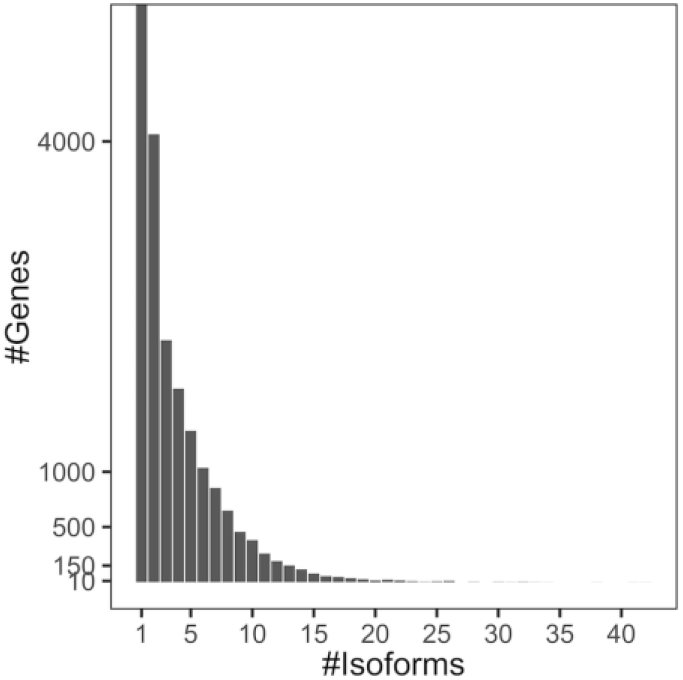
The distribution of the #genes according to the #annotated isoforms. The distribution of the number of genes for different number of annotated isoforms.

## Notes

### Competing Interest Statement

The authors have declared no competing interest.

### Summary of Updates

Added Salmon as one of the quantification methods benchmarked and updated all methods to current versions. The whole manuscript has been updated.

